# Perturbation of tonoplast sucrose transport alters carbohydrate utilization for seasonal growth and defense metabolism in coppiced poplar

**DOI:** 10.1101/2023.11.11.566720

**Authors:** Trevor T. Tuma, Batbayar Nyamdari, Chen Hsieh, Yen-Ho Chen, Scott A. Harding, Chung-Jui Tsai

**Author notes:** Corresponding author: Chung-Jui Tsai.

## Abstract

Non-structural carbohydrate reserves of stems and roots underpin overall tree fitness. These reserves also contribute to productivity under short-rotation management practices such as coppicing for bioenergy. While sucrose and starch comprise the predominant carbohydrate reserves of *Populus*, utilization is understood primarily in terms of starch turnover. The tonoplast sucrose transport protein SUT4 modulates sucrose export from source leaves to distant sinks during photoautotrophic growth, but the possibility of its involvement in remobilizing carbohydrates from sinks during heterotrophic growth has not been explored. Here, we used *PtaSUT4-*knockout mutants of *Populus tremula × P. alba* (INRA 717-1B4) in winter (cool) and summer (warm) glasshouse coppicing experiments to strain carbon demand and test for SUT4 involvement in reserve utilization. We show that epicormic bud emergence, subsequent growth, and xylem hexose increases were delayed or reduced in *sut4* mutants following lower temperature ‘winter’ coppicing. Depletion of stem reserves during post-coppice regrowth was not impaired in *sut4*, but *sut4* winter maintenance costs may have been higher in metabolic terms. Bark accrual of abundant defense metabolites, including salicinoids, chlorogenic acids, and flavonoid products, was prioritized in the summer, but attenuated in *sut4* mutants. Summer sprout growth was not reduced in *sut4* compared to controls. Together, our results point to shifting priorities for SUT4 modulation of seasonal trade-offs between growth and other priorities in the stem and emerging buds during reserve utilization in *Populus*.

## Introduction

Renewable energy production from feedstocks like tree crops is expected to help reduce fossil fuel use and greenhouse gas emissions responsible for global warming (Dhillon and von Wuehlisch 2013; Nieminen et al. 2012; Ragauskas et al. 2006). *Populus* species (e.g., poplar, aspen, cottonwood, and their respective hybrids) have proven useful for this purpose due to their fast growth, high biomass production, and ability to grow in marginal environments (Bryant et al. 2020; Chudy et al. 2019; Kauter et al. 2003). Autumn or winter coppicing, where trees are harvested near ground level and new shoots regenerate from the stump for repeated harvesting without replanting, is a common practice in short-rotation woody biomass production (Ek et al. 1983; Hytönen 1994). Feedstock biomass yields from coppicing can rival those of trees grown from seedlings or cuttings (Labrecque et al. 2023; Proe et al. 1999).

The success of coppicing is influenced by the utilization of non-structural carbohydrate reserves available from sink organs for new growth (Karp and Shield 2008; Luostarinen and Kauppi 2005). Sink carbohydrate reserves facilitate bud emergence and new shoot growth in the spring before source organ photosynthesis becomes active and support growth after defoliation or stem girdling events (Falk et al. 2020; Gleason and Ares 2004; Regier et al. 2010). In general, it is considered that both soluble sugars and starch comprise storage reserves, with exchange between the constituent pools occurring on daily and seasonal time-scales in trees (Martínez-Vilalta et al. 2016; Tixier et al. 2018). It is widely held that reserve dynamics are important to trade-offs between storage, growth, defense, and protection, all of which influence tree fitness (Blumstein et al. 2022).

Starch is regulated diurnally (Hartmann and Trumbore 2016; Klein and Hoch 2015; Noronha et al. 2018), and therefore has been the main focus of most studies on reserve carbohydrate utilization. However, significant diurnal fluctuations in leaf sucrose reserves occur in *Populus* (Wang et al. 2022; Zhang et al. 2014). Recent evidence suggests that in the absence of starch, other carbohydrates, presumably including sucrose, can provide the reserves needed to sustain new vegetative growth (Wang et al. 2022). Under extreme carbon limitation, carbohydrate storage can supersede growth as a priority (Huang et al. 2021; Piper and Fajardo 2016; Sala et al. 2012; Wang et al. 2022; Wiley et al. 2019). Under such conditions, trees may rely instead on alternative substrates such as lipids (Fischer et al. 2015; Huang et al. 2021) and hemicelluloses (Schädel et al. 2009) to sustain respiration and growth. It is worth noting that phenolic glycoside salicinoids, which are abundant defense compounds in *Populus* species, are turnover-resistant during severe carbon deprivation (Hillabrand et al. 2023). This highlights the importance of chemical defense in relation to growth and storage as a carbon destination in these species. Among the diverse metabolic and osmotic demands that potentially govern plant-wide competition for sucrose, limited attention has been given to the physiological importance of its intracellular compartmentalization.

Transmembrane sucrose transporters (SUTs) facilitate the movement of sucrose in all plants (Ayre 2011; Kühn et al. 1999). Several phylogenetically and subcellularly distinct isoforms are known (Peng et al. 2014). To date, one of the most explored proteins with a potential involvement in tree carbohydrate partitioning has been the tonoplast sucrose efflux transporter, SUT4 (Payyavula et al. 2011; Schneider et al. 2012). In *Populus*, *SUT4* is expressed at a relatively high level compared to most other *SUTs* throughout the plant (Payyavula et al. 2011). *SUT4* gene expression varies with a range of abiotic stresses including light, temperature, and drought (Xu et al. 2017). Effects of SUT4 impairment on turgor, osmotic adjustment, growth, and antioxidant metabolism in leaves of poplar suggest the potential for SUT4 to impact many homeostatic processes through sensing-signaling pathways that remain to be elucidated (Frost et al. 2012; Harding et al. 2020; Harding et al. 2022). Notably, *SUT4*-knockout (KO) impacts starch homeostasis, which suggests the potential for a conditional role of SUT4 in integrating utilization of the two major carbohydrate storage forms in *Populus* (Harding et al. 2022). Of special note for the present study, stem *SUT4* transcripts are detected at comparatively higher levels in the winter than in the summer of several temperate tree species including *Populus* (Dobbelstein et al. 2018; Ko et al. 2011; Sreedasyam et al. 2023).

These observations are all consistent with the general notion that SUT4 may be involved with the control of carbohydrate utilization in sink organs of *Populus*. Here, we used coppicing as a tool to investigate the role of tonoplast sucrose trafficking in sprouting under carbon-limiting conditions. A *SUT4-*KO effect on resprouting, osmolyte, or defense metabolite investments would imply that physiological trade-offs during reserve utilization are governed in part by vacuolar sucrose efflux. Two seasonal regimes were utilized in order to assess potential seasonal effects on SUT4 function. Possible contributions of SUT4 function in terms other than vacuolar sucrose efflux *per se* to ‘winter’ fitness are discussed. We also compared *SUT4*-KO effects on carbohydrate utilization under warm summer conditions where *sut4* growth was normal but defensive metabolite reserves were small.

## Materials and Methods

### Plant production and maintenance

*PtaSUT4*-KO mutants and the *Cas9* empty vector control lines used in this study were generated in the *P. tremula* × *P. alba* hybrid (INRA 717-IB4) by CRISPR/Cas9 mutagenesis as reported previously (Harding et al. 2022; Zhou et al. 2015) and grown in a glasshouse located in the Whitehall Forest at the University of Georgia. Supplemental LED lighting was used to maintain a 16-hour photoperiod, with photosynthetic photon flux density of at least 600 µmol m^-2^ s^-1^ at mid canopy. Plants were propagated by single-node cuttings as described (Frost et al. 2012) year-round for the experiments. Once established, rooted propagules were transferred to 1-gallon pots containing commercial soil mix (Fafard 3B) supplemented with Osmocote (15-9-12, NPK 6-month slow release) and maintained with daily watering until coppicing.

### Coppicing experiments

Several weeks prior to the winter experiment with post-coppice monitoring (December-January), twelve control (six wild-type [WT] and six *Cas9)* and twelve *sut4* mutants (four plants each of three independent events: KO-18, KO-25, and KO-51), were positioned with equal and random spacing on the floor bench of the glasshouse. From that point on, indoor high temperatures ranged from 20°C to 30°C after midday with long cool mornings, and nighttime temperatures routinely reaching 10°C. Ten days prior to coppicing, the number of fully expanded leaves was reduced to forty per plant to minimize differences in water uptake and photosynthetic capacity. Plants ∼1.5 m in height and approximately 4-months-old were then coppiced to ∼25 cm stumps and any remaining leaves were removed. Shoot tissues were collected for metabolic analysis and watering was reduced after coppicing to avoid root anoxia. Day length ranged from 10.2 hours to 10.7 hours. Supplemental heating prevented indoor glasshouse temperatures from dropping below 10°C. Freezing did not occur inside the glasshouse.

For the summer experiment with post-coppice monitoring (June-August), plants were managed as described above and were approximately the same height at coppicing as those in the winter experiment. Ten control plants (six WT and four *Cas9)* and twelve *sut4* mutants (four plants each of KO-18, KO-25, and KO-51 as above), were coppiced. Outdoor temperatures for the duration of the summer coppicing experiment ranged from 20°C to 33°C, with an average of 26°C. Day length ranged from 12.9 to 13.4 hours. Evaporative cooling prevented glasshouse temperatures from exceeding a high of 35°C during the day. Nighttime temperatures inside the glasshouse ranged from 15°C to 25°C. The coppicing experiments were carried out twice to ensure general reproducibility in post-coppice sprouting comparisons. Results from the most comprehensive of the two experiments are reported here (Figure S1).

### Tissue collection (pre- and post-coppicing)

Immediately after coppicing (t=0), bark and xylem tissues were collected from the basal end of the harvested stem (*n* = 8 – 12). At that time (t=0), xylem sap was also collected (described below). Additional bark and xylem tissues were collected from the stump tops when epicormic bud elongation was measured (t=1), and again during sprout extension (t=2). All tissues were sampled from the same individuals throughout the course of the experiments, snap frozen in liquid N and stored at -80°C. At t=0 and t=2, root samples (terminal 6 cm of soft elongating roots excluding tips) were washed in deionized water, dried, and snap frozen in liquid N. For starch, coarse root segments 4-6 mm in diameter were collected only at t=2.

### Xylem sap collection

Xylem sap was collected from the stumps immediately after coppicing following a previously published approach (Siebrecht and Tischner 1999). Bark was removed from the top 4 cm of the stump to expose the xylem cylinder and to avoid mixing of phloem and xylem sap during collection. The razor-cut surface of the wood was tamped clean with a damp kimwipe. The first 400 µl of xylem sap was discarded. Over the next 45 min, ∼100-200 µl of xylem sap was collected with a pipette from each stump and snap frozen in liquid nitrogen. Frozen sap was stored at -80°C until analysis.

### Bud emergence

Epicormic bud emergence was recorded daily beginning at t=0. The largest (dominant) epicormic bud that emerged after coppicing of each tree was photographed and its length determined at t=1 using the ImageJ software package (https://imagej.net/ij/index.html). Due to seasonal growth differences, sampling at times t=1 and t=2 was carried out 10 and 16 days, respectively, after summer coppicing and 11 and 20 days, respectively, after winter coppicing (Supplemental Figure S1). All tissue sampling occurred under sunny conditions between 11AM – 2PM.

### Biomass analysis from post-coppice sprouts

Effects of winter and summer conditions on new aerial biomass production after coppicing were measured in a third independent coppicing experiment. Plants (*n* = 6 – 11) ∼1.5 m in height were coppiced in winter and summer as described above. New sprouts were allowed to develop for ∼1 month after coppicing to determine biomass of shoots produced from stumps. Leaves (including petioles) and stem tissues of all sprouts were dried in a forced air oven for 48 hours at 60°C.

### Biomass analysis from vegetative propagules

In addition to the coppicing experiments, cohorts of ∼2 m control and *sut4* mutant plants (*n* = 7– 14 plants) grown from single internode stem cuttings using our standard vegetative propagation procedure were harvested in mid-summer and mid-winter. Plant height was measured at harvest, and leaves (including petiole), bark, and wood (de-barked stem) were collected to determine biomass allocation patterns during winter and summer growth. Root tissues were washed in deionized water and dried.

### Metabolite analysis

Tissues were freeze-dried using a FreezeZone 2.5 (Labconco), Wiley milled through a 40-mesh sieve (Thomas Scientific), and further ball-milled at 1500 rpm in a MiniG tissue homogenizer (Spex SamplePrep) to a fine powder. An aliquot of tissue powder (10.0 ± 0.5 mg, *n* = 8 – 12) was used for metabolite profiling by gas chromatography-mass spectrometry (GC-MS) as detailed in Harding et al. (2022). Adonitol (Sigma-Aldrich) was used as an internal standard. Samples were analyzed on an Agilent 7890A GC coupled to an Agilent 5975C single quadrupole MS detector (Agilent Technologies). Peaks were deconvoluted using AnalyzerPro (SpectralWorks) and putative identities were assigned based on the NIST08, the Agilent Fiehn metabolomics library, and an in-house library of authentic standards (Harding et al. 2022). A standard mix containing succinic acid, sucrose, glutamic acid, fructose, glucose, and ascorbic acid was loaded at the beginning and end of each sample set to monitor derivatization and instrument performance. Xylem sap samples (*n* = 8 – 12) were allowed to thaw completely on ice with mixing, and 10 μl were dried for derivatization and GCMS analysis as described above.

Another tissue powder aliquot (5.0 ± 0.5 mg, *n* = 8) was used for profiling by ultra-performance liquid chromatography coupled with quadrupole time-of-flight mass spectrometry (UPLC-QTOF). Tissue extraction was performed as above (Harding et al. 2022) with the exception that ^13^C_6_ –cinnamic acid and D_5_–benzoic acid were included as internal standards. Extracts were filtered through a 0.2 μm PTFE (polytetrafluoroethylene) filter (Agilent Technologies) and resolved on an Agilent Zorbax Eclipse Plus C18 column (2.1 × 50 mm, 1.8 μm) using an Agilent 1290 Infinity II UPLC coupled to an Agilent 6546 LC/Q-TOF tandem MS. The mobile solvent A comprised water and 0.1% formic acid and mobile solvent B comprised acetonitrile with 0.1% formic acid at a flow rate of 0.5 mL min^-1^. The elution gradient was 3% B from 0 – 0.5 min, linear gradient to 15% B over 3.5 min, isocratic at 30% B over 4.5 min, linear gradient to 50% B over 5.5 min, and then to 100% B over 7 min, followed by a 3-min post-column run. Column temperature was 45°C. MS data were acquired in negative mode with the following parameters: gas temperature, 250°C; nebulizer gas, 40 psi; nozzle voltage, 500 V; and capillary voltage, 400 V. Data were processed using the MassHunter software suite Version 11.0 (Agilent). Peak identities were confirmed using an in-house library of authentic standards, when available, or assigned by searching against an accurate-mass NIST20 library. The highest database match score (usually >85%) obtained was used to support the putative identity of the QTOF metabolites.

GC-MS and LC-QTOF peak intensities were normalized by their respective internal standards and sample dry weight, with the exception of tissue sucrose and hexose (fructose and glucose) levels which are reported as percent dry weight based on their respective calibration curves. Together, sucrose and the secondary metabolite salicortin comprise approximately two-thirds of the solvent-extractable metabolite load in leaves and bark of the experimental poplar clone, with hexoses comprising half of the remainder (Harding et al. 2020; Harding et al. 2022). In light of their smaller contribution to overall metabolite abundance, the remaining lower abundance constituents were compared using log2-transformed ratio based on normalized peak areas.

### Starch

Starch content was determined by enzymatic digestion with α-amylase and amyloglucosidase (Chow and Landhäusser 2004). In brief, 10 mg of lyophilized, Wiley- and ball-milled tissue powder (*n* = 8 – 12) was pre-extracted three times with 1 ml of methanol by vortexing and sonication. The tissue pellet was resuspended in 0.5 ml of buffer (0.1 M sodium acetate, pH 5.0, with 5 mM CaCl_2_) containing 500 U of α-amylase (Sigma A4551) and incubated for 2 hr at 65°C. Samples were cooled and 10 µl of sodium acetate buffer (pH 5.0) containing 5 U amyloglucosidase (Sigma A1602) was added for further digestion at 50°C for 48 hr with shaking. Digests were then diluted with 500 µl of H_2_O, vortexed, centrifuged, and 20 μl of the diluted supernatant were dried and derivatized for GC-MS as described above. Chromatograms contained only glucose and adonitol internal standard peaks. Total starch content was calculated based on a glucose standard curve and expressed as % dry weight.

### Condensed tannins (CTs)

Freeze dried tissue powder (10 mg, *n* = 8) was extracted in 600 µl of methanol for 15 minutes in a sonicator bath to extract interfering chlorophyll. The depigmented pellet was dried and CT content was estimated using the butanol-HCl method (Porter et al. 1986) as described (Harding et al. 2022). CT concentration was determined based on absorbance at 550 nm (SpectraMax M2; Molecular Devices) using purified aspen leaf CTs as standards.

### Chlorophyll content

Ten mg of freeze-dried powder (*n* = 8 – 12) was extracted twice in 750 µl of cold 90% acetone – 10% water, with sonication and centrifugation. Absorbances at 645 nm and 663 nm were used to determine chlorophyll *a*, chlorophyll *b*, and total chlorophyll content (µg g^-1^ DW) as described by Lichtenthaler (1987).

### Statistics

Univariate repeated measures analysis of variance (ANOVA) was used to determine significant differences at each stage of coppicing re-growth over time, between genotypes, and their interactions using *R* v4.1.0 in Rstudio (version 2022.12.353). All other reported genotypic contrasts were assessed via Student’s *t-*test. Linear regression was used to test for the relationships between sucrose or starch depletion and bud elongation of control (WT and *Cas9* lines) or *sut4* mutants between t=0 and t=1 of the winter and summer coppicing experiments. Statistical significance was determined at a predetermined alpha level (*e.g.*, 0.05).

## Results

### Seasonal coppicing response of sut4 mutants

The rate at which new epicormic buds appeared on stumps after coppicing was largely similar between genotypes under the summer glasshouse conditions (Figure 1A, C). The total biomass of new shoots at t=2 in the *sut4* mutants was also similar to the controls despite initially faster bud elongation in *sut4* mutants (t=1) during the summer (Figure 1E-F). In contrast, epicormic bud emergence was delayed by several days in *sut4* mutants compared to controls in the winter (Figure 1B, D). In addition, bud elongation at t=1 was slower and total biomass at t=2 was reduced in the *sut4* mutants compared to control plants following winter coppicing (Figure 1E-F). Biomass yield and partitioning of more advanced coppice sprouts 32 and 40 days after summer and winter coppicing, respectively, were measured in separate plant cohorts (Table 1).

**FIGURE 1:**
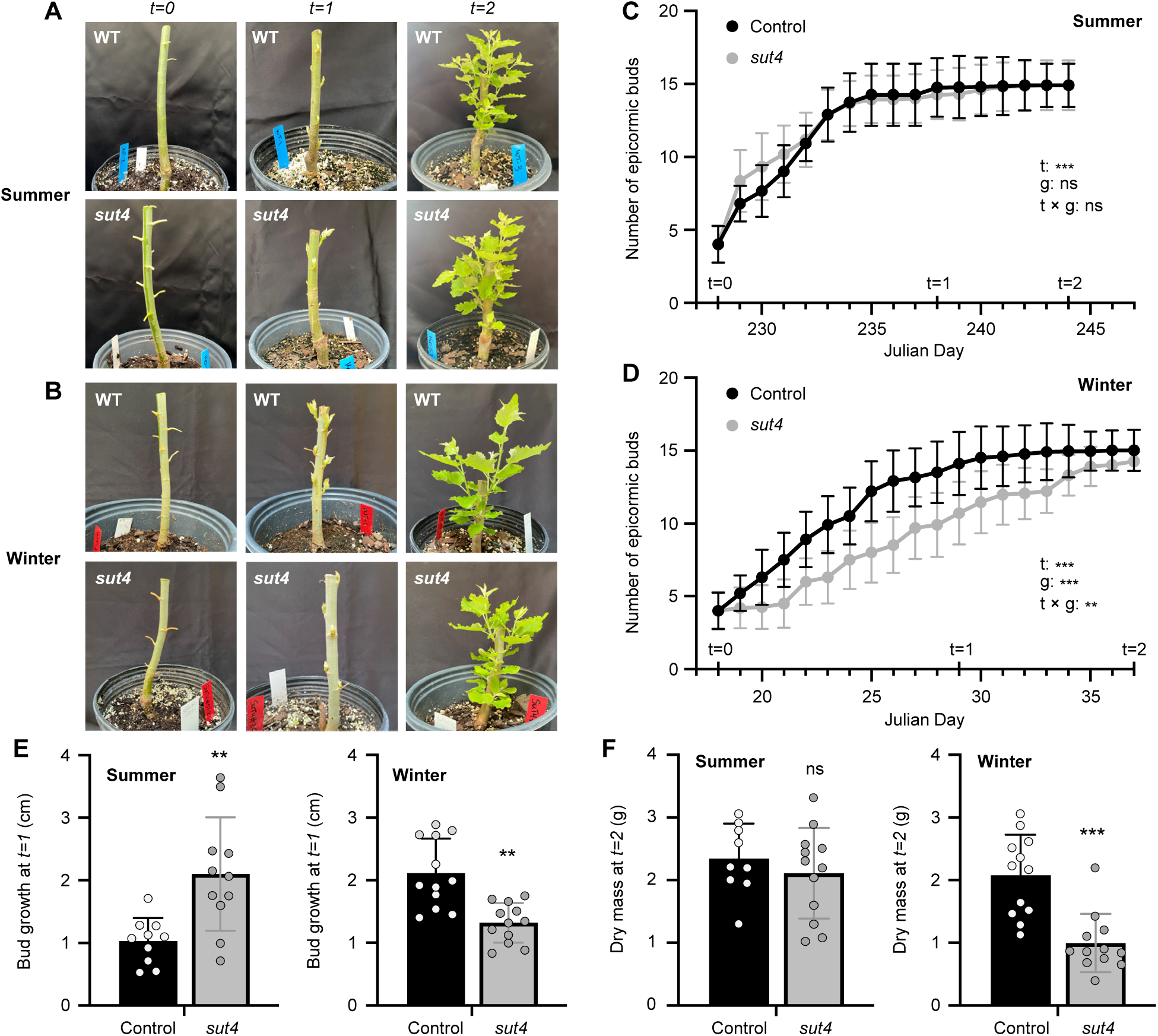
Epicormic bud emergence and growth following coppicing. (A-B) Representative images of *sut4* and control poplar stools under summer (A) and winter (B) conditions immediately after coppicing (t= 0), during epicormic bud expansion (t=1), and during sprout growth (t=2). (C-D) Bud emergence after summer (C) and winter (D) coppicing. Black and grey represent control and *sut4* plants, respectively. Data are mean ± SD of *n* = 8 – 12 biological replicates. The effects of time (t), genotype (g), or their interaction (t × g) were assessed by a repeated measures two-way ANOVA. (E) Length of largest (dominant) epicormic bud at t=1. Statistical significance was determined by Student’s *t* test between control and *sut4* plants. (F) Sprout biomass at t=2. Statistical significance was determined by Student’s *t* test between control and *sut4* plants. ns, nonsignificant; *, *P* ≤ 0.05; **, *P* ≤ 0.01; *\*\*\*, P* ≤ 0.001.

Significantly more sprout leaf and stem dry mass were produced from *sut4* than control stumps under summer conditions (Table 1). Sprout biomass trended lower, but not significantly, in *sut4* than in control plants in the winter. Single shoots grown for 10-12 weeks from single internode stem cuttings rather than from coppiced stumps were slightly larger for control plants than *sut4* plants as has previously been reported for *SUT4-*silenced or KO plants (Frost et al. 2012; Harding et al. 2020; Payyavula et al. 2011), regardless of season (Table S1). We conclude that slight negative *sut4* effects on growth are the general rule, and that the strong biomass performance of *sut4* summer coppice sprouts (Table 1) may have a basis in metabolic findings described later.

### Sink carbohydrate reserves

Sucrose levels of basal stem tissues and roots (sinks) were generally higher in *sut4* than control lines at the time of coppicing (t=0) regardless of the season (Figure 2A). Sucrose levels decreased after coppicing in all stump sinks including roots in both genotypes. The most striking observation was that for xylem but not for other sinks, sucrose levels at t=0 and t=1were much higher in winter than in summer. This was especially notable for *sut4* (Figure 2A). In addition, the net reduction of xylem sucrose by t=2 following winter coppicing was greater in *sut4* than control stumps. Root sucrose was not as depleted as in bark or xylem by coppicing (Figure 2A).

**FIGURE 2:**
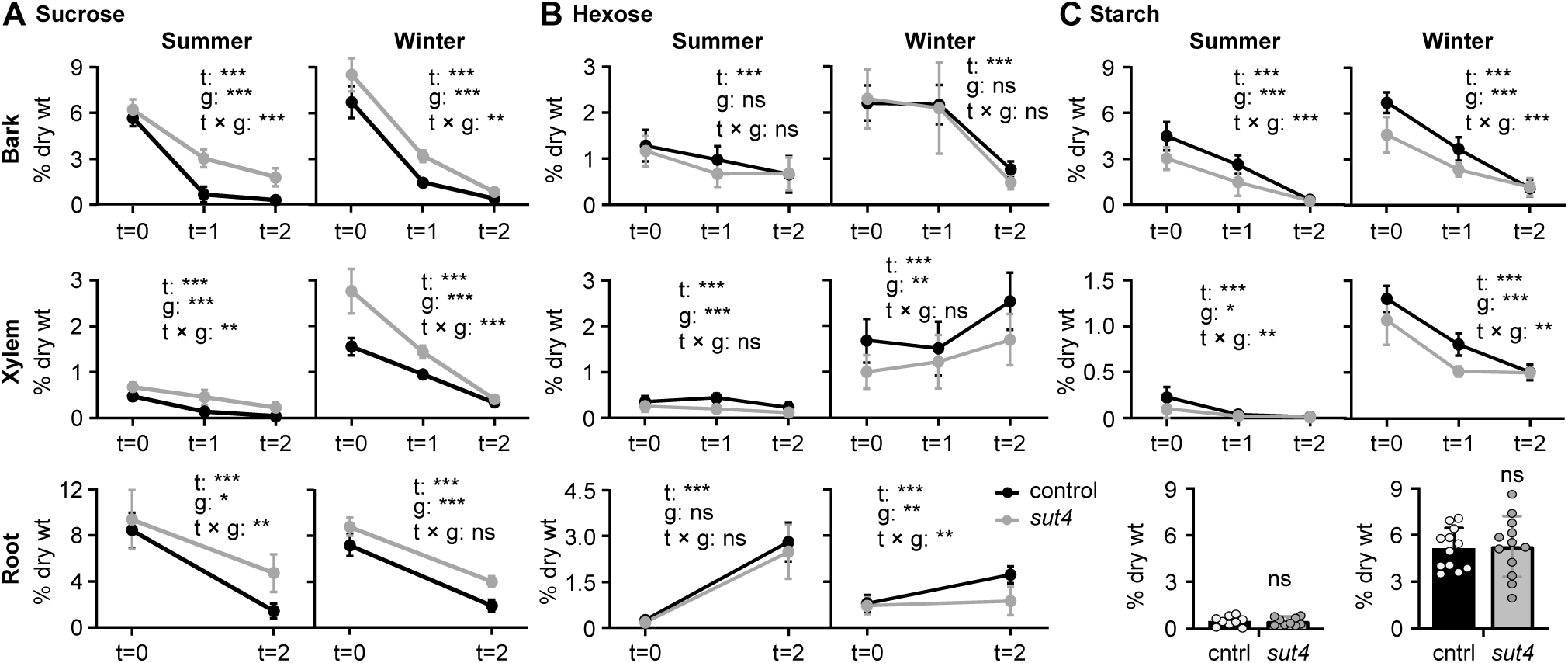
Non-structural carbohydrate abundance dynamics in stem and root tissues following coppicing. (A) Sucrose, (B) Hexose (sum of fructose and glucose), and (C) Starch. Black and grey are for control and *sut4* plants, respectively. Values are mean ± SD of *n* = 8 – 12 biological replicates. The effects of time (t), genotype (g) or their interaction (t × g) were assessed by repeated measures two-way ANOVA. Statistical significance for root starch levels at t=2 was determined by Student’s *t* test between control and *sut4* plants. ns, nonsignificant; *, *P* ≤ 0.05; *\*\*, P* ≤ 0.01; *\*\*\*, P* ≤ 0.001.

Bark hexose (sum of fructose and glucose) levels and trends differed little overall between genotypes, being 2-fold higher in winter than summer at t=0 but decreasing much more precipitously by t=2 in winter than in summer (Figure 2B). Similar to the trend with sucrose, xylem hexose levels were several-fold greater in winter than in summer at t=0. Unlike the case for sucrose, xylem hexose levels were higher in control than *sut4* mutants in dry mass terms. While xylem sucrose trended down after winter coppicing, xylem hexose trended up (Figure 2B). The average summer xylem hexose differential between control and *sut4* was 0.15% dry weight (versus an average content of ∼0.27% dry weight), and the winter xylem hexose differential was 0.6% dry weight (versus an average hexose content of 1.6% dry weight). Comparing the winter xylem sucrose and hexose data plots, sucrose/hexose ratios were consistently and substantially lower in control than *sut4*, in line with a *sut4* restriction on sucrose availability for winter transport and metabolic cleavage in the stem. Root hexose levels increased sharply in both genotypes after summer coppicing, but the increase was observed only in control plants after winter coppicing (Figure 2B).

Starch levels in the bark and xylem were higher in control than *sut4* plants in both the summer and winter, and remained so as starch decreased through t=1 (Figure 2C). Bark and xylem starch was depleted to similarly low levels in both genotypes by t=2, suggesting that starch abundance there had reached a minimum. Root starch content was similar in both genotypes at t=2 but was much less abundant in the summer than the winter (Figure 2C). Together, the data suggest a depletion of non-structural carbohydrates to less than 25% of pre-coppice levels in shoot organs. Root sucrose also decreased sharply, but starch was only measured at final harvest (t=2).

### Sucrose and starch utilization for bud growth

Slower post-coppice sprout emergence and growth in *sut4* than control trees in the winter led us to test whether sucrose and/or starch depletion between t=0 and t=1 in the bark predicted bud length at t=1 (Figure 3). Bud size did not correlate with sucrose depletion in the summer for either genotype (*sut4: R^2^ =* 0.17, *p* = 0.21, control*: R^2^ =* 0.06, *p* = 0.49), but the slope trended positive for *sut4* and negative for control plants (Figure 3A). Under winter conditions, bud length correlated positively and significantly with sucrose depletion in the *sut4* mutant trees (*R^2^ =* 0.39, *r=* 0.62, *p* = 0.03) but not in the controls (*R^2^ =* 0.16, *p* = 0.19) where in fact the slope trended negative as in summer (Figure 3A). Bud expansion did not correlate with changes in starch content under summer (*sut4: R^2^ =* 0.02, *p* = 0.68, control*: R^2^ =* 0.12, *p* = 0.439) or winter (*sut4: R^2^ =* 0.00, *p* = 0.99, control*: R^2^ =* 0.04, *p* = 0.54) conditions (Figure 3B). Overall, post-coppice bud elongation in *sut4* mutant, but not in control plants, tended to correlate positively with sucrose but not with starch depletion. Additional metabolic outcomes (to be described) pertinent specifically to inhibition of sucrose transport by SUT4 may have contributed to the improved significance for *sut4* (Figure 3).

**FIGURE 3:**
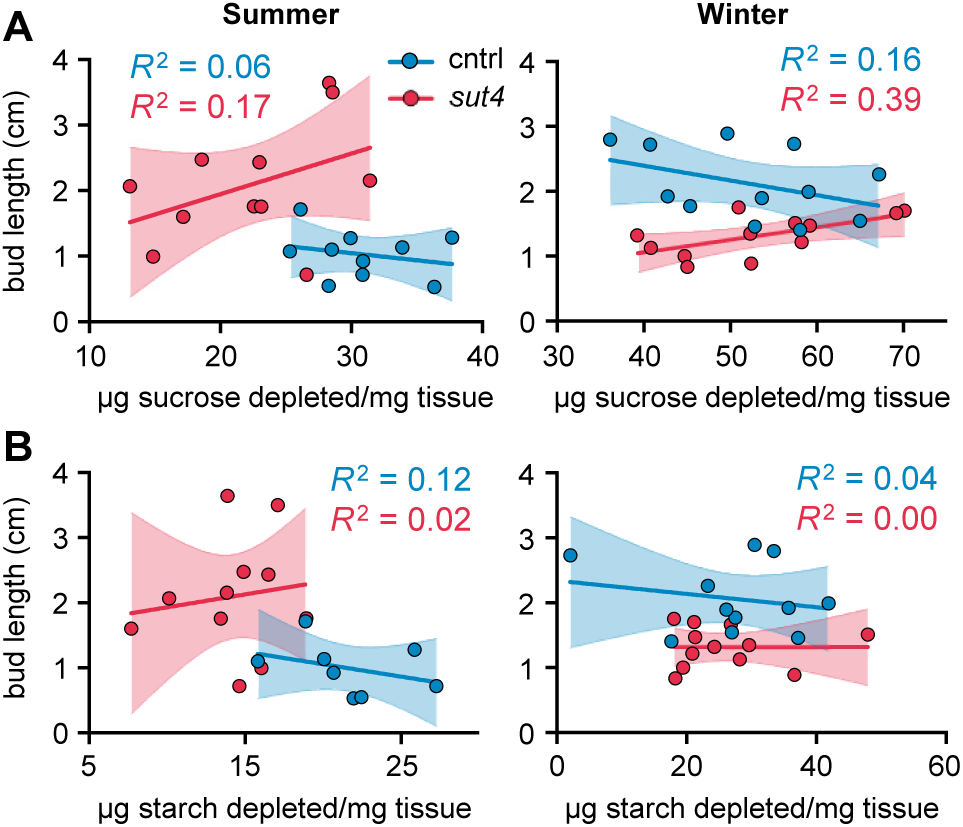
Epicormic bud length as a function of bark carbohydrate depletion under summer and winter conditions. (A-B) Bud length at t=1 versus sucrose (A) or starch (B) depletion between t=0 and t=1. Linear regression trendlines are depicted in blue and red for control and *sut4*, respectively (*n* = 8 – 12). Shading highlights the 95% confidence intervals.

### Xylem sap and root metabolites

Xylem sap sucrose abundance at t=0 was greater in winter than summer, but differed little between genotypes during either season (Figure 4A). Sap hexose levels did not differ between genotypes in the summer but were lower in *sut4* than control plants in the winter (Figure 4B). Xylem sap is also the route by which most root-assimilated nitrogen travels to the transpiring leaves (Siebrecht and Tischner 1999). Considering that nitrogen assimilation depends on carbon skeletons within the assimilating organ, Krebs cycle intermediate and amino acid contents in the xylem sap could reveal seasonal and genotypic differences in the roots’ systemically accessible carbon supply. Krebs cycle intermediates malate and citrate comprised most of the xylem sap metabolic load in both summer and winter, and were more abundant in *sut4* than control xylem saps (Figure 4C-D). Glutamine and glutamate were the most abundant amino acids we detected in xylem sap. Both trended higher in *sut4* than control xylem sap in the summer but were significantly lower in *sut4* than control sap in the winter (Figure 4C-D). These four metabolites were also among the most abundant in root tissues, but only malate mimicked the xylem sap patterns, being most abundant in *sut4* (Supplemental Figure S2).

**FIGURE 4:**
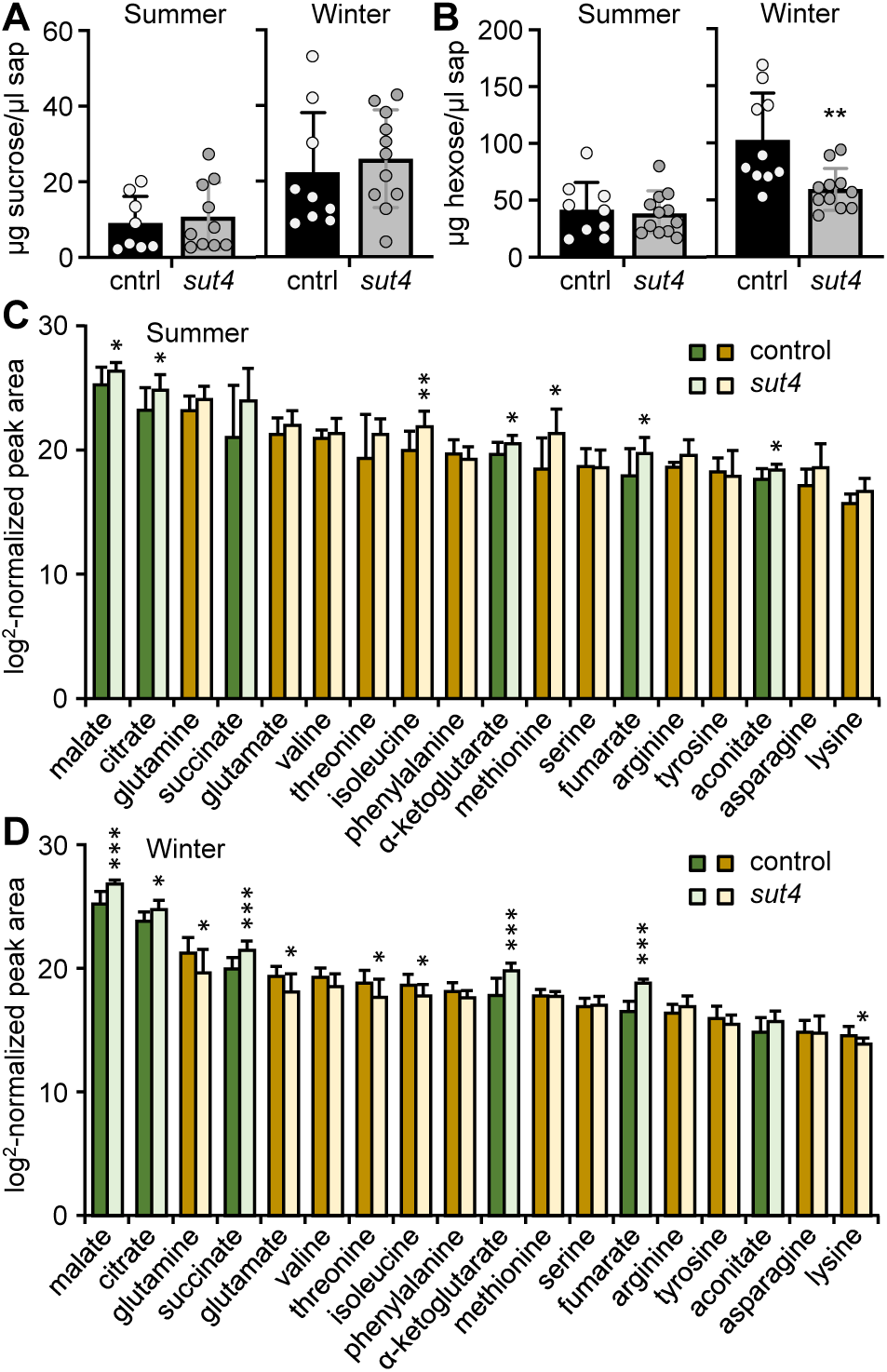
Xylem sap metabolites. (A) Sucrose, (B) Hexoses, (C-D) Abundance ranking of the most prevalent metabolites after sucrose and hexose in control and *sut4* sap in summer (C) and winter (D). Krebs cycle intermediates are in green and amino acids are in brown colors, with control and *sut4* in darker and lighter shading, respectively. Values represent the mean ± SD (*n* = 8-12). Statistical significance between genotypes was determined by Student’s *t* test. *, *P* ≤ 0.05; *\*\*, P* ≤ 0.01; *\*\*\*, P* ≤ 0.001.

### Organic acid metabolism in bark and xylem

Bark and xylem metabolite trends at t=0 roughly paralleled those observed in xylem saps, with amino acids being lower and Krebs cycle intermediates such as malic and citric acid higher in *sut4* mutants, most markedly in winter xylem (Figure 5). The Krebs cycle differentials increased during bud expansion, but the amino acid differentials changed relatively little in bark and not at all in xylem (t=1 and t=2) (Figure 5). Gluconic acid, which is associated with the pentose-phosphate pathway (Horecker 2002), exhibited a trend similar to that of the major Krebs cycle intermediates (Figure 5). Phenylpropanoid pathway precursors (e.g., quinic acid and shikimic acid) and intermediates (e.g., hydroxycinnamates and catechin) were detected at comparatively low levels in *sut4* bark and xylem throughout the course of both seasons.

**FIGURE 5:**
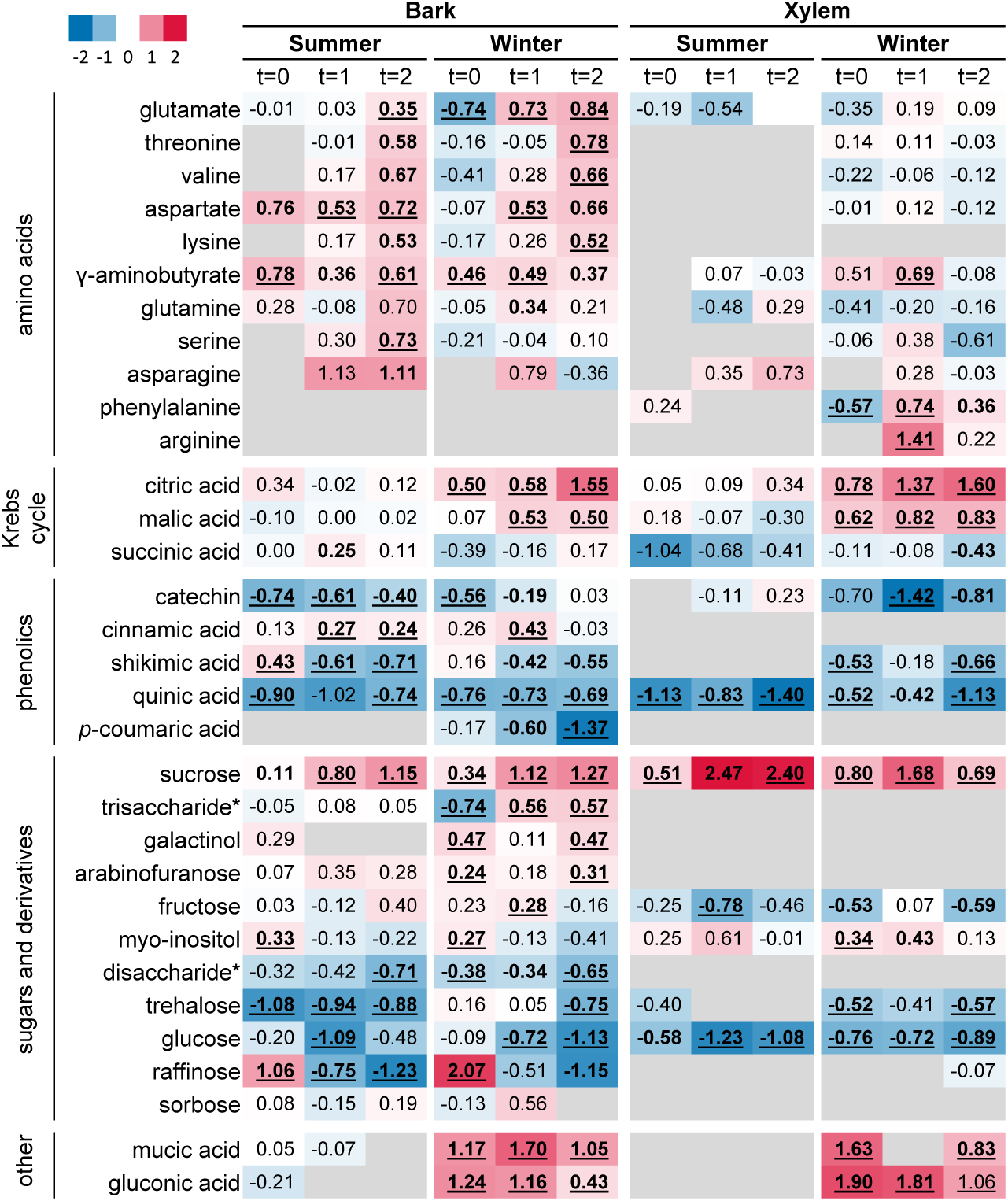
Post coppice metabolic reprogramming in bark and xylem tissues. Heatmap depiction of log_2_-transformed metabolite response ratio (*sut4*/control) in bark (A) and xylem (B). Blue and red colors indicate decreased or increased metabolite abundance in mutants, while grey indicates not detected. Significance between genotypes was obtained by Student’s *t*-test and is indicated in bold underlined (*P* ≤ 0.01) or bold font (*P* ≤ 0.05). Asterisks denote putative di- and tr-saccharides.

### Raffinose

Levels of the phloem-transported and well-known stress carbohydrate raffinose were greater in the bark of *sut4* plants at the time of coppicing (t=0), but much lower than in controls by t=2 (Figure 5). Raffinose precursors including myoinositol and galactinol exhibited less clear differentials.

### Phytochemical reserves for protection and defense

Provisioning of nonstructural sinks related to poplar defense has been associated with carbohydrate transport (Arnold et al. 2004; Kleiner et al. 1999). In *Populus* and *Salix* species, chlorogenic acids, salicinoids (e.g., salicortin and tremulacin), and condensed tannins (CTs) derived from quinate-shikimate-phenylpropanoid pathway are present in especially large abundance for fitness and stress tolerance (Harding et al. 2005; Tsai et al. 2006; Zhang et al. 2018). Chlorogenic acids (caffeoyl-quinate isomers) exhibited lower abundance in *sut4* relative to control bark at all time points in both the summer and winter coppicing experiments (Figure 6A). Salicortin, by far the most abundant defense metabolite, was detected at higher levels in the summer than the winter (Figure 6B). Bark salicortin abundance was substantially lower in the *sut4* mutants than control during the summer but did not differ between genotypes during the winter (Figure 6B). The closely related and less abundant tremulacin was also significantly lower in *sut4* mutant bark in the summer, but not the winter (Figure 6C). Bark CT levels were lower in *sut4* than control plants, especially in the summer (Figure 6D).

**FIGURE 6:**
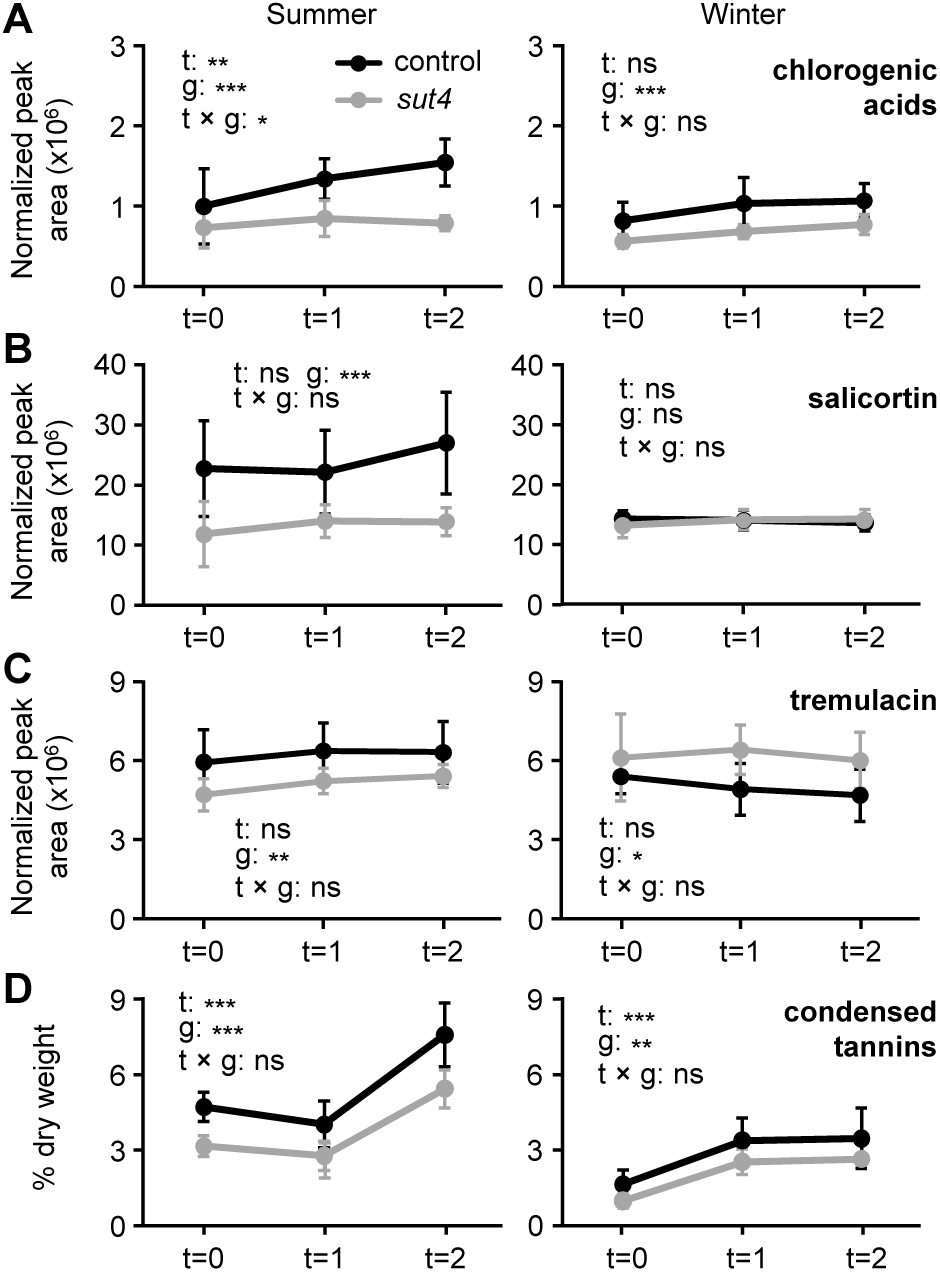
Bark secondary metabolites (A) Chlorogenic acids (sum of caffeoyl-quinate isomers), (B) salicortin, (C) tremulacin, and (D) condensed tannins. Values represent the mean ± SD of *n* = 8 biological replicates. The effects of time (t), genotype (g) or their interaction were assessed by repeated measures two-way ANOVA. *, *P* ≤ 0.05; *\*\*, P* ≤ 0.01; *\*\*\*, P* ≤ 0.001.

### Chlorophyll content

We examined bark chlorophyll contents as a proxy for photosynthetic capacity after coppicing. Bark chlorophyll content differed little between genotypes and remained relatively stable during summer coppicing experiment, but showed a transient decrease at t=1 during winter (Figure 7).

**FIGURE 7:**
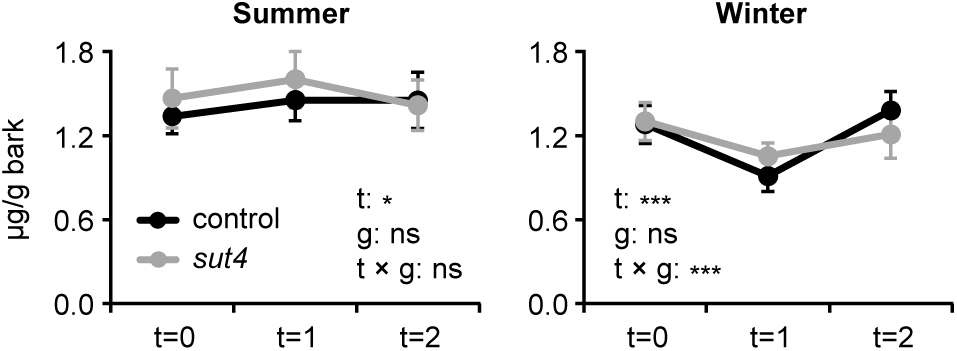
Bark chlorophyll content. Changes in bark chlorophyll following summer (A) and winter (B) coppicing. Black and grey are for control and *sut4* plants, respectively. Data are presented as the mean ± SD (*n* = 8-12). The effects of time (t), genotype (g) or their interaction were assessed by repeated measures two-way ANOVA. *, *P* ≤ 0.05; *\*\*\*, P* ≤ 0.001.

## Discussion

Trees produce nonstructural carbohydrate reserves to sustain respiration and osmoregulation for winter protection and spring bud flush during perennial growth (Furze et al. 2019). Therefore, and in light of growing environmental challenges due to climate change, plasticity of reserve utilization is likely to be important for species survival and ecosystem composition in the future (Blumstein et al. 2022; Furze et al. 2019; Martínez-Vilalta et al. 2016; Sevanto et al. 2014). One outcome of climate change is increasingly milder winters in temperate latitudes, and with that, changes in the stresses experienced by overwintering trees (Contosta et al. 2019). In light of this context and in consideration of agricultural practices such as fall or winter coppicing of trees, the studies presented here aimed to provide insights into the involvement of one of the better studied sucrose transporters of *Populus* in reserve carbohydrate remobilization. Our findings underscore the role of SUT4 when sprout growth depends on carbohydrate remobilization and delivery from stem reserves. Ultimately, *sut4* sprouts recovered from much of their initial growth deficit after several weeks of photoautotrophic growth (Table 1). Considering the temperature and osmotic sensitivity of *SUT4* gene expression (Xu et al. 2017), penalties and recovery are environmentally responsive processes subject to global climate change.

### Winter osmotic adjustments were compromised in xylem of sut4 mutants

Previous analysis of carbohydrate active gene expression in overwintering stem tissue of poplar supports involvement of starch degradation as well as of sucrose synthesis and trafficking during this stressful period (Ko et al. 2011). According to multiple studies involving temperate tree species, transcript levels for *SUT4* encoding the tonoplast sucrose transporter peak in the winter in both stem and apical buds (Figure S3) (Dobbelstein et al. 2018; Ko et al. 2011; Sreedasyam et al. 2023). Partial or full loss of SUT4 function in *Populus* has been associated with compromised carbohydrate metabolism, turgor and osmotic responses to drought (Frost et al. 2012; Harding et al. 2020; Harding et al. 2022). Given the association of desiccation and altered turgor with winter freezing, we propose in light of our findings under warm versus cool growing conditions that SUT4-mediated sucrose trafficking within stems has anticipatory roles in winter fitness, irrespective of the occurrence of ice crystal formation. Although freezing temperatures were not applied in this study, sugar osmolyte differences between summer and “winter” glasshouse conditions were observed at t=0 in xylem (Figure 2 A-B, Figure 5). We suggest that these results are consistent with an anticipation of more injurious chilling, triggered in our experiments by overnight cooling followed by slow morning warming of roots in relation to shoots, and seasonal differences in light quality.

### Starch did not compensate for the negative sut4 effect on “winter” sprouting

Under the cool-temperature “winter” glasshouse conditions in this study, stump reserves of starch and sucrose were severely, but similarly reduced in control and mutant lines (Figure 2). Yet, despite ample starch reserves, epicormic bud emergence and perhaps also subsequent expansion were markedly slower in *sut4* mutants (Figure 1D-E). In a contrast, post-coppice reserve depletions in summer were smallest in *sut4* by t=1 (Figure 2A-B), but epicormic bud expansion was markedly slower in the control plants. In fact, early epicormic bud growth of controls was nominally slower in summer than winter (Figure 1E). This will be discussed later in connection with summer versus winter defensive metabolite allocations in bark (Figure 6).

While early growth did not correlate with starch depletion in either genotype regardless of season, early growth was positively associated with bark sucrose depletion only in *sut4* and only in winter (Figure 3). It is probable that sucrose trafficking both within (buds) and between organs (bark to bud) normally increases in importance during the winter in line with higher *SUT4* gene expression in winter stems and buds (Figure S3) (Dobbelstein et al. 2018; Ko et al. 2011; Sreedasyam et al. 2023). Nevertheless, the “winter” limitation to *sut4* sprout initiation may have been most acute when competition by xylem sinks that are normally supported by increased efflux of vacuolar sucrose, was greatest. If the apparent greater dependence on sucrose in the *sut4* mutants (Figure 3) suggested a greater sucrose limitation, then the rate of sucrose depletion should have been lower in the bark and xylem between winter t=0 and t=1 in *sut4* relative to the controls. However, no such difference was evident (Figure 2A). Therefore, we considered the possibility of an additional metabolic cost associated with *sut4,* particularly in winter sucrose utilization, as indicated by the results of our regression analyses (Figure 3).

### Winter organic acid production and sprout growth in sut4

Krebs cycle intermediates malic and citric acid exhibited relatively higher levels in winter *sut4* bark and xylem (Figure 5), suggesting a potential increase in respiratory loss of metabolic carbon for cellular maintenance at the expense of biogenesis for growth. This conclusion is consistent with the observation that the abundance differentials for growth-related amino acids whose carbon skeletons are derived from the Krebs cycle did not increase as much (Figure 5). Why other Krebs cycle functions, including respiration, would be preferentially stimulated by perturbed vacuolar sucrose efflux in the winter than in the summer, remains unclear. However, the altered osmotic dynamics in winter may have led to higher maintenance costs in *sut4*.

Furthermore, gluconic acid levels were also significantly elevated in *sut4,* indicating increased activity in pentose phosphate pathway activity in winter. This could be attributed to DNA repair, NADPH requirements for fatty acid synthesis for membrane repair, and reactive oxygen species control (Horecker 2002; Jiang et al. 2022). The greater abundance of mucic acid (Figure 5), an oxidized form of galactose also known as galactaric acid, in the *sut4* plants during winter may indicate an increase in carbohydrate oxidation despite increased fluxes through essential antioxidant and maintenance pathways.

It is also plausible that the greater accrual of Krebs cycle organic acids in *sut4* contributed to cytosolic pH buffering. This buffering may have been necessitated due to reduced proton efflux from the vacuole since SUT4 is a proton-sucrose efflux symporter (Ayre 2011). Utilization of sucrose that is not exported from the vacuole usually depends at least in part on the activity of vacuolar invertase (Koch 2004; Ruan 2014; Schulz et al. 2011). As is the case for *SUT4*, invertase expression is modulated by stresses such as drought and cold (Chen et al. 2015; Roitsch and González 2004; Xue et al. 2016). The efflux of invertase product monosaccharides (hexose) from the vacuole by proton symporters would in principle support cytosolic pH buffering by a SUT4-like mechanism. However, hexose efflux from vacuoles is considered to be largely passive, with other mechanism such as coupled symport and antiport mechanisms coming to light more recently (Slewinski 2011; Zhu et al. 2021).

### Stress amelioration by SUT4

Sugar osmolytes are known for their established roles in scavenging reactive oxygen species, enhancing cold tolerance, and aiding in embolism refilling during winter (Charrier et al. 2018; Tarkowski and Van den Ende 2015). The abundant expression of cell wall invertases in poplar winter stems supports the notion that hexoses may be involved in these winter processes (Figure S3) (Ko et al. 2011; Sreedasyam et al. 2023). Besides the overall reduction in hexose, sharp post-coppice reductions in other important sucrose-derived protective osmolytes like raffinose between t=0 and t=2 were observed in *sut4* compared to control bark (Figure 5). Impairment of drought induced increases in leaf raffinose have also been attributed to *SUT4* inhibition by RNAi (Frost et al. 2012). Interestingly, prior to coppicing (t=0), bark raffinose was much more abundant in *sut4* than control bark (Figure 5). In other work carried out under relatively cool conditions similar to those of the present study, leaf raffinose was higher in the *SUT4*-impaired lines along with evidence for cytosolic sucrose increases versus the controls (Harding et al. 2022). Together these studies suggest a beneficial effect of raffinose under drought stress or in response to the threat of chilling stress in *Populus*, but they also point toward a role for SUT4 in this process under increasingly carbon-limiting conditions when heterotrophic reserves are being depleted.

The elevated *SUT4* expression in winter stem (Ko et al. 2011; Sreedasyam et al. 2023) (Figure S3) may be influenced by factors other than temperature, such as abiotic stresses (i.e., desiccation) known to increase *SUT4* expression (Xu et al. 2017; Xue et al. 2016). Interestingly in this regard, drought stress increased endogenous *SUT4* expression in various organs of *SUT4*-RNAi *Populus*, as well as in other dicot species where leaf expression has been reported (Xu et al. 2017; Xue et al. 2016). These findings raise interesting questions with regard to how increased frequencies of unseasonal episodes of biotic (e.g., insect or fungal) or abiotic (e.g., drought or heat) stresses due to transient warming might affect SUT4 and the timing of reserve utilization for growth and protection in overwintering trees.

### SUT4 and seasonal metabolic priorities

In summer, the metabolic priority of post-coppice control stumps seemed to shift toward much higher accruals of resource-intensive defense metabolites such as chlorogenic acids, salicortin, and CTs than in winter (Figure 6). Interestingly, summer defense metabolite levels including those of salicortin were substantially lower in *sut4* than control lines (Figure 6B). At the same time, *sut4* sprout biomass production equaled or exceeded, at least transiently, that of the controls during summer but trailed that of controls under the winter conditions (Figure 1 and Table 1). Salicortin abundance in bark and leaves of the poplar hybrid used in this study can rival those of sucrose, and maintenance costs of salicortin may be high or low depending on how turnover is measured (Harding et al. 2020; Kleiner et al. 1999; Ruuhola and Julkunen-Tiitto 2000). Therefore, depending on the magnitude and developmental timing of metabolic costs of salicinoids in very young, rapidly expanding organs, phytochemical investment in summer control bark may have been sufficiently large to metabolically neutralize the sprout biomass production advantage of control stumps seen in winter when defense metabolite investment was lower (Figure 1E-F).

Turgor control may also be considered as a contributing factor to the relative improvement in *sut4* sprout growth under summer conditions. Turgor of fully expanded leaves is greater in *sut4* than control plants because of higher sugar osmolyte levels (Harding et al. 2020; Harding et al. 2022). This implicates a role for *SUT4* in modulating early expansion growth in control plants where *SUT4* expression increases as leaves transition from sink to source organs (Payyavula et al. 2011). The increase in *SUT4* expression would support sucrose efflux into the cytosolic compartment for metabolite biosynthesis (e.g. salicortin and CTs) at the expense of rapid turgor accrual. The present findings suggest a complex developmental dynamic influenced by light, temperature, nutrient and source:sink ratio in which SUT4 has a role. *Populus* species maintain constitutive levels of salicinoids even under severe carbon limitation (Fabisch et al. 2019; Hillabrand et al. 2023), and our results may suggest an environmentally conditioned impact of vacuolar sucrose sequestration on the competition between growth and this important phytochemical sink.

In conclusion, *Populus* SUT4 was revealed through seasonal differences in the coppicing responses of KO mutant lines to have a role in nonstructural carbohydrate allocations for growth and winter protection. Future research may investigate the effects of *SUT4*-KO under field conditions to further explore seasonal influences on SUT4 function and perennial growth habit.

## Data and Materials Availability

Data and material will be made available upon reasonable request.

## Supplemental data

**Table S1.** Biomass allocation of mature control and *sut4* poplar.

**Figure S1.** Schematics of coppicing experiments.

**Figure S2.** Root metabolites.

**Figure S3.** Seasonal variation of *SUT* and representative carbohydrate gene expression.

## Acknowledgements

The authors thank Gilles Pilate of INRA, France for providing poplar clone INRA 717-1B4, Suzzanne Tate for greenhouse assistance, Khadijeh Mozaffari for LC-QTOF assistance, and W. Patrick Bewg, Samantha Surber, Amya Covington, McKenzie Drowns, Hazel Quarterman, Chaitanyi Pasupuleti, and H. Hansini Fernando for assistance with sampling and tissue processing.

## Authors’ Contributions

T.T.T, S.A.H, & C.J.T. conceived the study and designed all experiments. T.T.T. performed all experiments and analyzed the data. T.T.T, B.N., & S.A.H, performed metabolic analyses. Y.H.C. and C.H. helped with expression analysis. T.T.T. prepared the draft manuscript, which was revised by S.A.H. with contributions from C.J.T.

