## Supplementary material for "Perturbation of tonoplast sucrose transport alters carbohydrate utilization for seasonal growth and defense metabolism in coppiced poplar": Table S1 (from another version of this preprint: version 1, 2023-11-14)

**TABLE S1:** Biomass allocation of mature control and *sut4* poplar.

| | Summer | | Winter | |
|---|---|---|---|---|
| | Control | *sut4* | Control | *sut4* |
| Height (cm) | 208.27 ± 13.97 | 197.14 ± 10.63* | 194.71 ± 6.78 | 182.33 ± 8.44** |
| No. of leaves | 45.27 ± 2.37 | 48 ± 3.16* | 41.71 ± 1.80 | 45.33 ± 2.24** |
| Leaf mass (g) | 30.54 ± 3.56 | 34.68 ± 7.20 | 26.66 ± 3.25 | 34.33 ± 3.33*** |
| Bark mass (g) | 10.17 ± 2.47 | 9.91 ± 1.47 | 9.07 ± 1.89 | 9.58 ± 1.61 |
| Wood mass (g) | 36.38 ± 7.46 | 30.63 ± 4.95* | 37.62 ± 6.90 | 29.28 ± 4.33* |
| Stem (bark+wood) mass (g) | 46.55 ± 9.73 | 40.54 ± 6.23 | 46.69 ± 6.30 | 38.87 ± 4.20** |
| Root mass (g) | 12.02 ± 2.51 | 10.16 ± 2.31 | 11.65 ± 2.31 | 9.68 ± 1.31* |

Values represent mean and SD of $n$ = 7 – 14 biological replicates. Student's *t*-test was used to determine statistical significance between genotypes. *, $P \leq 0.05$; **, $P \leq 0.01$; ***, $P \leq 0.001$.

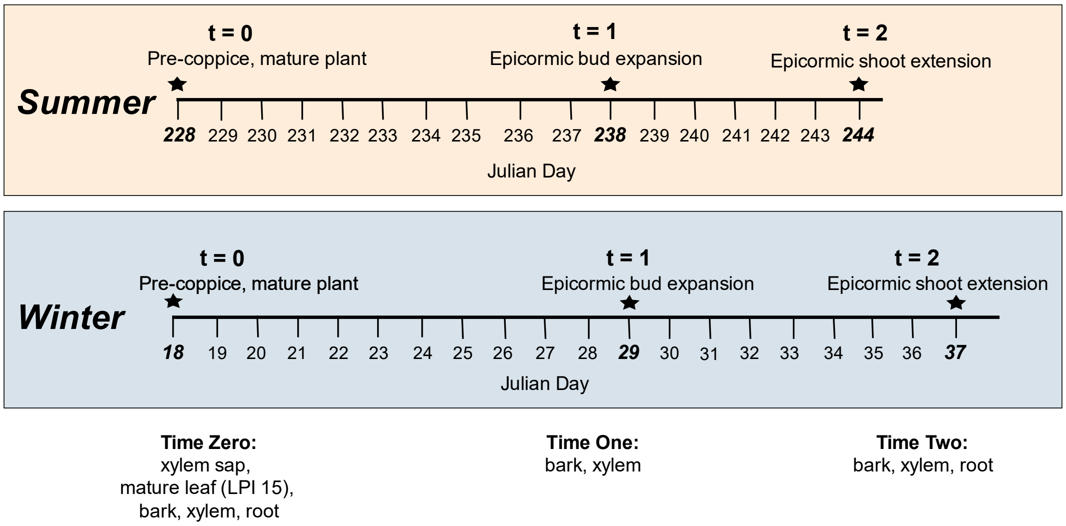


**Figure S1.** Schematics of coppicing experiments.
Bark and xylem were sampled from the coppiced stool t=0, 1, 2, with t=0 corresponding to the date of coppicing. For epicormic bud expansion measurements, the length of the largest (dominant) epicormic bud on each stool at t=1 was obtained by Image J software. Days until post-coppice sampling at t=1 and 2 differed between the winter and summer experiments in order to normalize for seasonal effects on bud emergence and sprout growth rates.

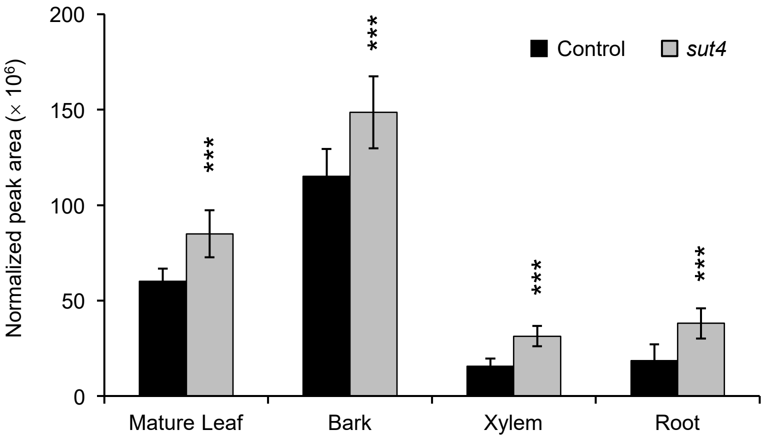


**Figure S2.** Sucrose levels in mature tree organs.
Data represent mean ± SD of n = 14-19 biological replicates. Control and *sut4* mutants are shown in black and grey, respectively. ***, $P \leq 0.001$ according to Student's *t* test.

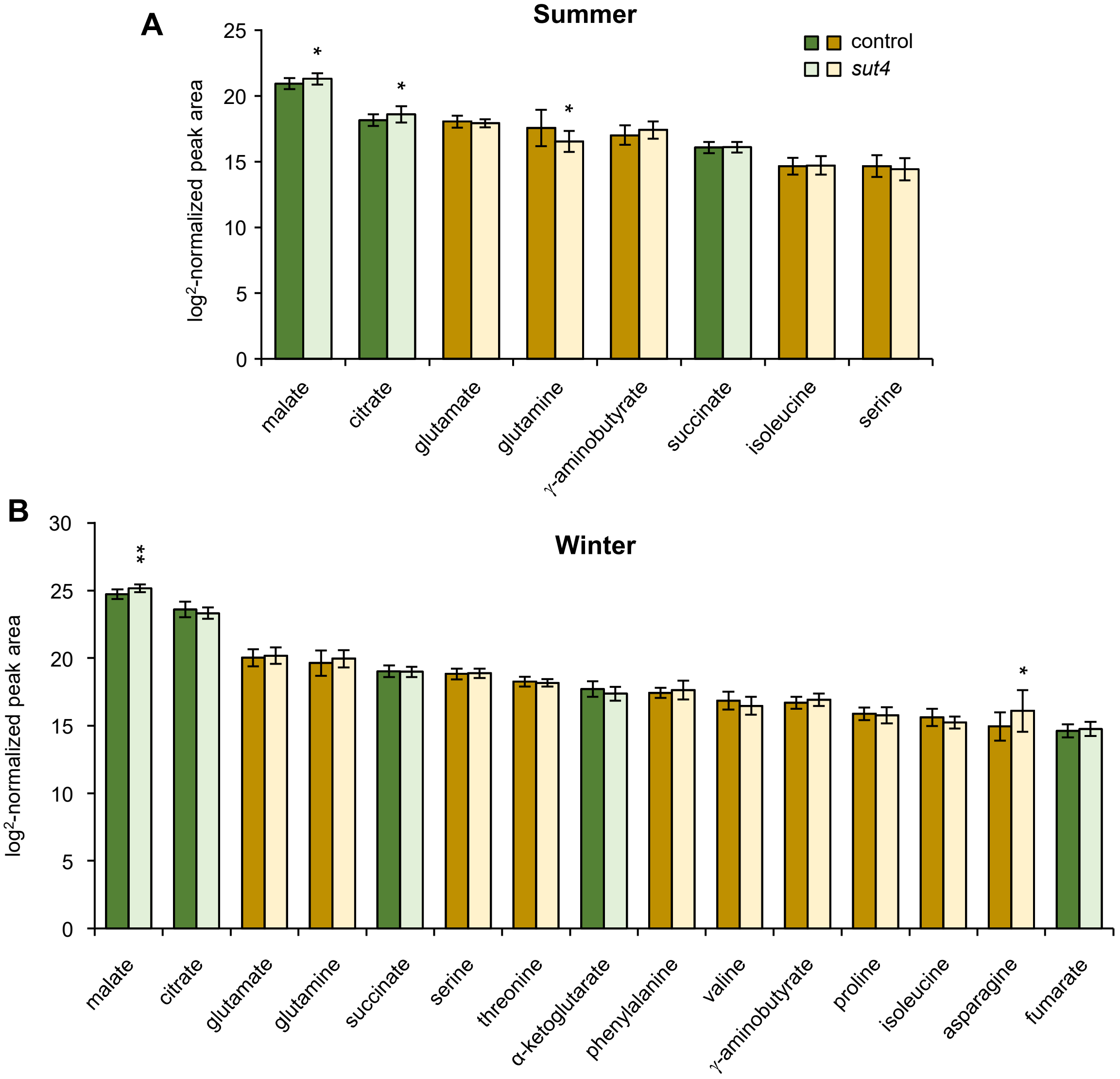


**Figure S3.** Root metabolites.
Normalized and $\log_2$ transformed peak areas of steady state Krebs cycle intermediates (green colors) and amino acids (brown colors) in control (darker shading) and *sut4* (lighter shading) plants in summer (A) and winter (B). Data are mean ± SD (n = 7-14 control or 13-19 mutant plants). Statistical significance was determined by Student's *t* test between control and *sut4* plants ($P \leq 0.01$; *, $P \leq 0.05$).

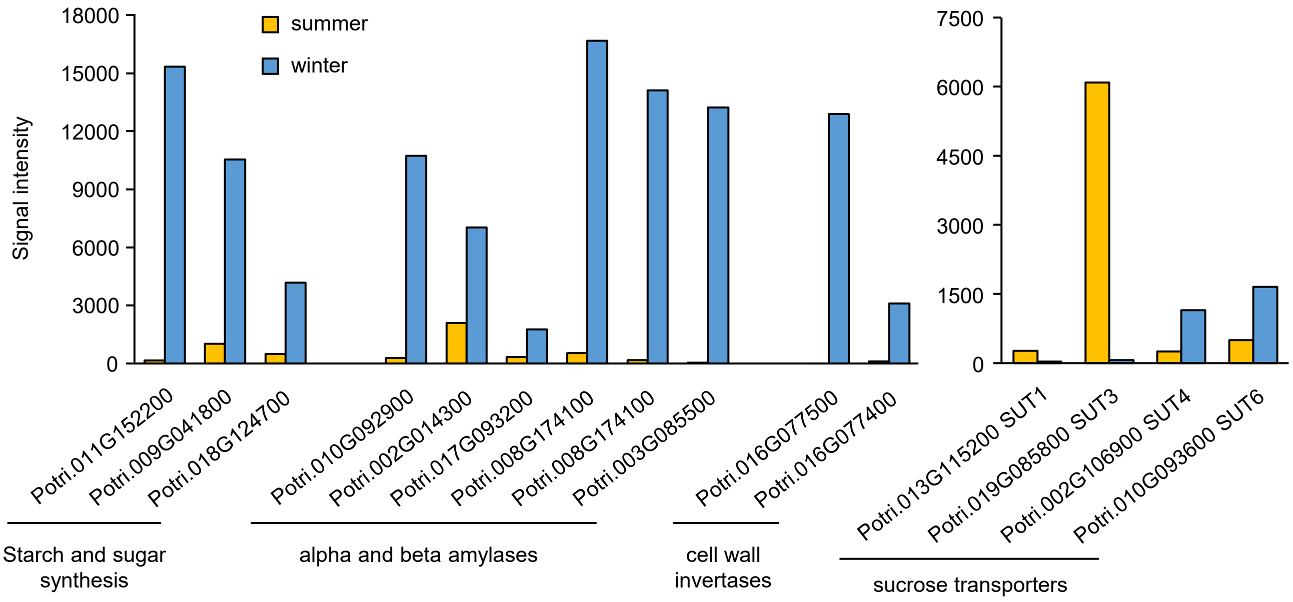


**Figure S4.** Seasonal variation of SUT and representative carbohydrate gene expression.
Expression of representative genes involved in carbohydrate metabolism in summer or winter *Populus* stem tissues. Data were obtained from Affymetrix microarray hybridization of Ko et al. (2011) based on annotation of Xue et al. (2015).


Ko, J.-H., Prassinos, C., Keathley, D., Han, K.-H. and Li, C. (2011) Novel aspects of transcriptional regulation in the winter survival and maintenance mechanism of poplar. Tree Physiology, 31, 208-225.

Xue L-J, Alabady MS, Mohebbi M, Tsai C-J (2015) Exploiting genome variation to improve next-generation sequencing data analysis and genome editing efficiency in *Populus tremula* × *alba* 717-1B4.  Tree Genetics and Genomes 11: 82.
